# Intrinsically disordered N-terminal domain (NTD) of p53 interacts with mitochondrial PTP regulator Cyclophilin D

**DOI:** 10.1101/2021.07.23.453429

**Authors:** Jing Zhao, Xinyue Liu, Alan Blayney, Yumeng Zhang, Lauren Gandy, Fuming Zhang, Robert J. Linhardt, Jianhan Chen, Christopher Baines, Stewart N. Loh, Chunyu Wang

## Abstract

Mitochondrial permeability transition pore (mPTP) plays crucial roles in cell death in a variety of diseases, including ischemia/reperfusion injury in heart attack and stroke, neurodegenerative conditions, and cancer. To date, cyclophilin D is the only confirmed component of mPTP. Under stress, p53 can translocate into mitochondria and interact with CypD, triggering necrosis and cell growth arrest. However, the molecular details of p53/CypD interaction are still poorly understood. Previously, several studies reported that p53 interacts with CypD through its DNA-binding domain (DBD). However, using surface plasmon resonance (SPR), we found that full-length p53 (FLp53) binds to CypD with *K_D_* of ~1 μM, while both NTD-DBD and NTD bind to CypD at ~10 μM *K_D_* (Fig. 1C and 1D). Thus, instead of DBD, NTD is the major CypD binding site on p53. NMR titration and MD simulation revealed that NTD binds CypD with broad and dynamic interfaces dominated by electrostatic interactions. NTD 20-70 was further identified as the minimal binding region for CypD interaction, and two NTD fragments, D1 (residues 22-44) and D2 (58-70), can each bind CypD with mM affinity. Our detailed biophysical characterization of the dynamic interface between NTD and CypD provides novel insights on the p53-dependent mPTP opening and drug discovery targeting NTD/CypD interface in diseases.

## Introduction

The tumor suppressor p53 plays a central role as a hub protein regulating diverse cellular processes, including but not limited to cell cycle arrest, DNA repair, apoptosis, necrosis, autophagy, and senescence^1^. p53 can not only mediate apoptosis in a transcription-dependent manner, but also use transcription-independent mechanisms^2,3^. For example, p53 can translocate to the mitochondrial surface, where it physically interacts with both anti- and pro-apoptotic Bcl-2 family members to inhibit or activate their respective functions, regulating mitochondrial outer membrane permeabilization and apoptosis^4^. Under cellular stress, p53 was reported to accumulate in the mitochondrial matrix and trigger mitochondrial permeability transition pore (mPTP) opening^5^. A physical interaction of p53 with the mPTP regulator cyclophilin D (CypD) in mitochondria was detected in oxidative stress/salinomycin/dexamethasone/cDAPK1-induced necrosis^5–8^ and cell growth arrest during tumorigenesis^9^. A robust p53–CypD complex forms during mouse brain ischemia reperfusion injury (stroke model) and ischemia-reperfusion-induced acute kidney injury^5,10^. Reduced p53 levels^5^ or pretreatment with cyclosporin A^10^ prevented the formation of this complex. At present, CypD is the only established component of mPTP^11^. Thus, mitochondrial CypD-p53 interaction likely plays an important role in the regulation of mitochondria-mediated necrosis and apoptosis; however, the structural mechanism of CypD-p53 interaction is not well understood.

p53 comprises multiple domains including the intrinsically disordered N-terminal domain (NTD), the well-folded DNA-binding domain (DBD), the tetramerization (TET) domain, and the intrinsically disordered C-terminal regulatory domain (CTD) (Fig. 1A). The nuclear localization signal sequences (NLS) were located between DBD and TET, which mediate the migration of p53 into the cell nucleus. NTD is composed of two transactivation sub-domains (TAD1 and TAD2) and the proline rich region (PRR). The intrinsically disordered nature of NTD enables specific interactions with multiple partners and further supports versatile p53 functions^12–15^. The DBD binds to target DNA at promoter regions and initiates the transcription of genes, which contains most of the oncogenic mutations of p53^16,17^. The DBD was reported to be also responsible for interactions of p53 with Bcl-2 family proteins and CypD^4,5,18,19^, while the role of NTD is less studied. Interestingly, a naturally occurring isoform, p53Ψ (27KD), which is the product of an alternatively splicing event and lacks the major portions of the DBD, the NLS, and the TET of p53, can translocate to mitochondria and mediate the mPTP opening through a direct interaction with CypD^20^. Like p53Ψ, TP53 exon-6 truncating mutations, which occurs at higher than expected frequencies in human tumors, also promotes tumorigenesis in a transcriptional independent manner through binding to CypD^21^. p53Ψ and exon-6 p53 truncating mutants (such as R196* and R213*, truncation in the middle of DBD), lack an intact DBD; but they can still interact with CypD, leading to increased mPTP permeability^20,21^ and pro-metastatic phenotype. This evidence raises the question whether p53 interacts with CypD at sites other than DBD.

**Fig. 1.**
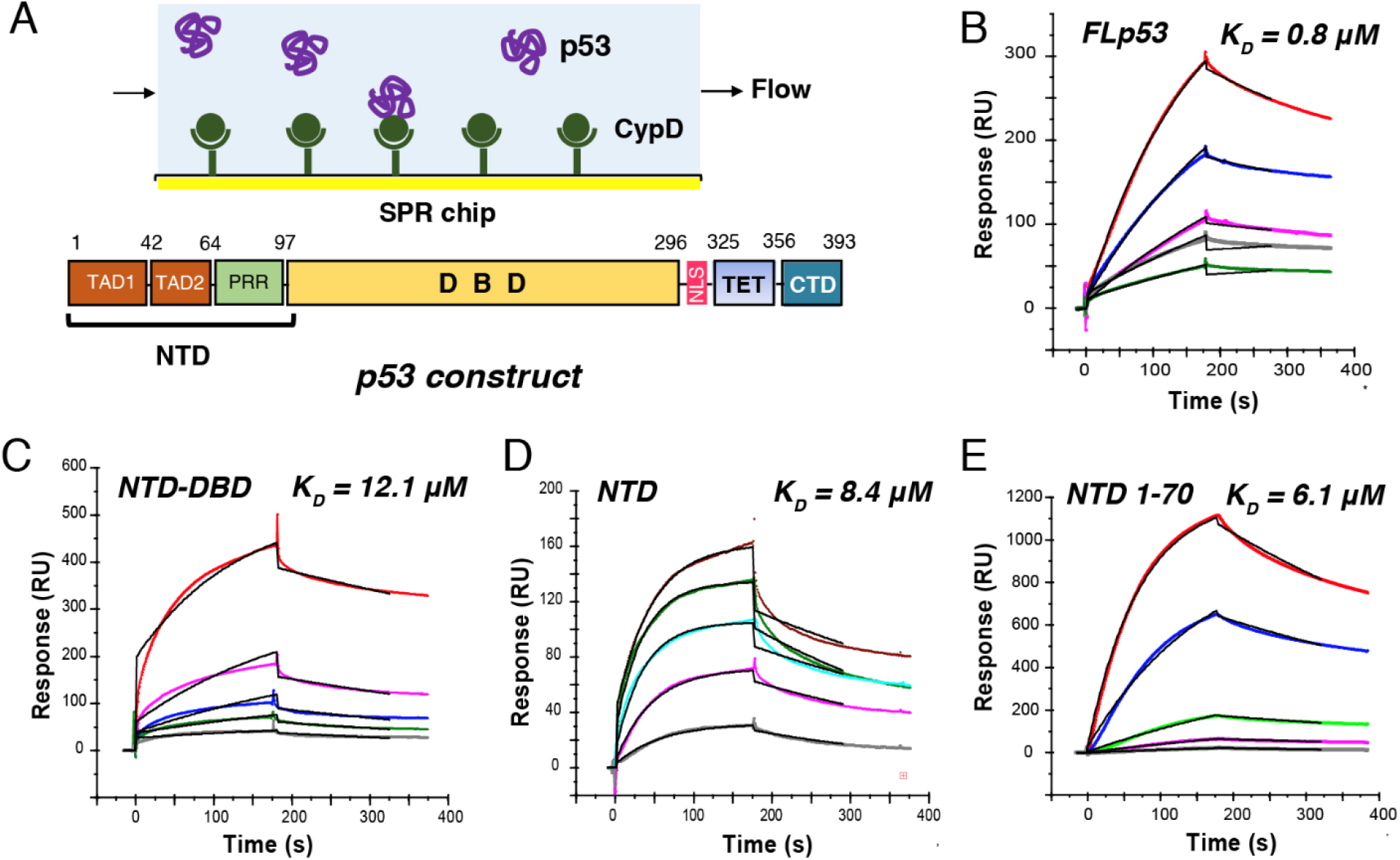
p53 binds to CypD at μM affinity determined by surface plasmon resonance (SPR). (A) SPR scheme (top) and p53 construct (bottom). (B) SPR sensorgrams of the full length p53 (FLp53)-CypD interaction. The concentrations of FLp53 (from top to bottom) were 2.0, 1.0, 0.5, 0.25, and 0.13 μM, respectively. (C) SPR sensorgrams of the NTD-DBD-CypD interaction. The concentrations of NTD-DBD (from top to bottom) were 50, 25, 12.5, 6.25, and 3.13 μM, respectively. (D) SPR sensorgrams of the NTD-CypD interaction. The concentrations of NTD (from top to bottom) were 40, 25, 12.5, 6.25, and 3.13 μM, respectively. (E) SPR sensorgrams of the interaction between CypD and NTD1-70. The concentrations of NTD1-70 (from top to bottom) were 10, 5, 2.5, 1.25, and 0.63 μM, respectively. All SPR sensorgrams are fitted using 1:1 Langmuir binding model (Black curves) in BIAevaluation software 4.0.1.

In this study, utilizing surface plasmon resonance (SPR) and nuclear magnetic resonance spectroscopy (NMR), we have shown for first time that the intrinsically disordered N-terminal domain (NTD) of p53 is the main binding site of CypD, instead of the DBD. NMR and MD simulation showed a dynamic and electrostatic interface between NTD and CypD. The binding interface on p53 was further narrowed to NTD 20-70. These biophysical data shed new light to CypD-p53 interaction with important implications for drug discovery targeting this interface in multiple diseases.

## Results

### p53 binds to CypD through N-terminal domain (NTD)

To investigate the role of p53 domains in p53-CypD interaction, different p53 constructs were tested for binding to an SPR sensor chip immobilized with CypD (Fig. 1A). As shown in Fig 1B, full-length p53 (FLp53) binds to CypD with a *K_D_* of ~1 μM. Both NTD-DBD and NTD bind to CypD at ~10 μM *K_D_* (Fig. 1C and 1D), indicating that, instead of DBD, NTD is the major CypD binding site on p53. The difference in binding affinity can be attributed to the different oligomerization states, as FLp53 exists primarily as a dimer^22^ and NTD and NTD-DBD are present mainly in monomeric form (data not shown). CypD is a mitochondrial matrix proline isomerase whose enzymatic activity is essential to triggering mPTP opening^23^. To further investigate if the proline rich region (PRR) in NTD is involved in CypD binding, a C-terminal truncated mutant NTD 1-70 was generated. NTD 1-70 interacts with CypD with a *K_D_* of ~6 μM (Fig. 1E). This result demonstrates that the PRR of NTD is not required for the p53-CypD interaction, suggesting that TAD1 and TAD2 are likely the major binding sites. This has also been seen in other protein-protein binding interactions of p53, such as with the coactivator binding domain of the CREB binding protein^24^. Vaseva et al. previously showed that p53 residues 80–220 is the region required for CypD interaction by pull-down experiments^5^. In a subsequent study, Lebedev et al. showed that CypD causes aggregation of both FLp53 and isolated DBD *in vitro*^18^. In contrast, here we identified by SPR that NTD 1-70 is the most crucial p53 region for CypD interaction.

### NTD mimics FLp53 binding to CypD

To further characterize the interaction, we compared the chemical shift perturbations (CSP) of isotopically labeled CypD upon the addition of FLp53, NTD and NTD-DBD using ^1^H–^15^N heteronuclear single-quantum correlation (HSQC) NMR spectroscopy. Since NMR chemical shifts are sensitive to changes in the chemical environment, CSPs can be used to identify the binding interface. Upon the addition of FLp53 (1: 0.2), HSQC spectrum of CypD experienced significant peak movement (Fig. 2C). Residue S77 exhibited the largest CSP, followed by A12 and G80. Other residues with significant CSPs include F83, G65, Q111, K76, K15, and A101. The addition of more NTD to a ratio of 1: 0.3 (CypD: FLp53) also caused significant CSPs in residues S77, F83, G65, and A101 (Fig. 2A). CSPs induced by FLp53 and NTD were superimposed in Fig. 3D (top). CypD CSPs in the region around S77 and G80 are similar for NTD and FLp53, while less binding was found in region around A12 for NTD (Fig. D). Notably, NTD caused larger CSPs in residues A101 and C115 than FLp53, indicating that NTD may access more binding sites on CypD. In contrast, the addition of NTD-DBD (1: 0.3) did not cause significant CSPs in residues S77 and F83, which are the dominant binding sites for FLp53 (Fig. 3B). Superimposition of CSPs induced by FLp53 and NTD-DBD (Fig. 3D, bottom) show different patterns. This result indicates that NTD alone is a better mimic of FLp53 binding to CypD than NTD-DBD. This phenomenon may be due to a masking effect of DBD on NTD, as a recent study showed that NTD has direct but weak interactions with DBD in FLp53, which affects DNA-binding activity^13^. Thus, an enhanced shielding effect could be observed for NTD-DBD. Previous pull-down experiments had shown that CypD interacts with p53 residues 80–220, corresponding to the N-terminal portion of the DBD^5^, while our investigation suggests that NTD mimics most of the FLp53 binding to CypD. In previous NMR work by Lebedev et al., much smaller perturbations were observed on CypD spectrum upon the addition of DBD at a relatively high ratio (1:2 CypD:DBD), indicating very weak binding for DBD. Here we show that at a much lower CypD to p53 ratio (1:0.2), FLp53 causes significant CSPs, most prominently around residue S77. NTD mimics this binding pattern around S77 at a similar ratio (1:0.3) which is absent in the titration of NTD-DBD (Fig. 2D), suggesting NTD to be a more important domain for CypD interaction than NTD.

**Fig. 2.**
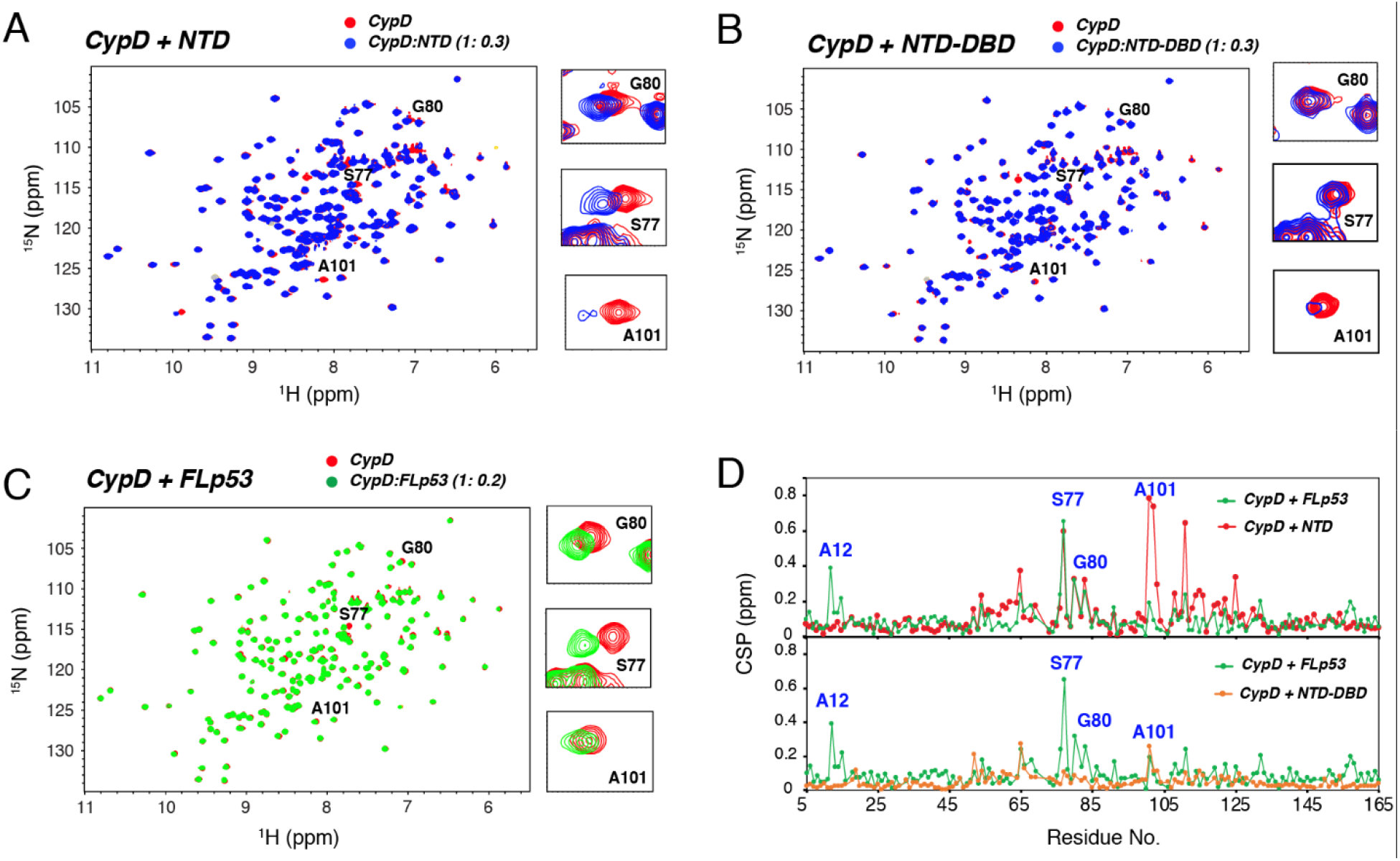
NTD mimics full-length p53 binding to CypD better than NTD-DBD. (A) ^1^H-^15^N HSQC spectra of CypD upon the addition of NTD (1: 0.3). (B) ^1^H-^15^N HSQC spectra of CypD upon the addition of NTD-DBD (1: 0.3). (C) ^1^H-^15^N HSQC spectra of CypD upon the addition of FLp53 (1: 0.2). Residues with large CSPs were magnified on the right. (D) CSPs cause by FLp53 superimposed onto NTD (top) and NTD-DBD (bottom).

**Fig. 3.**
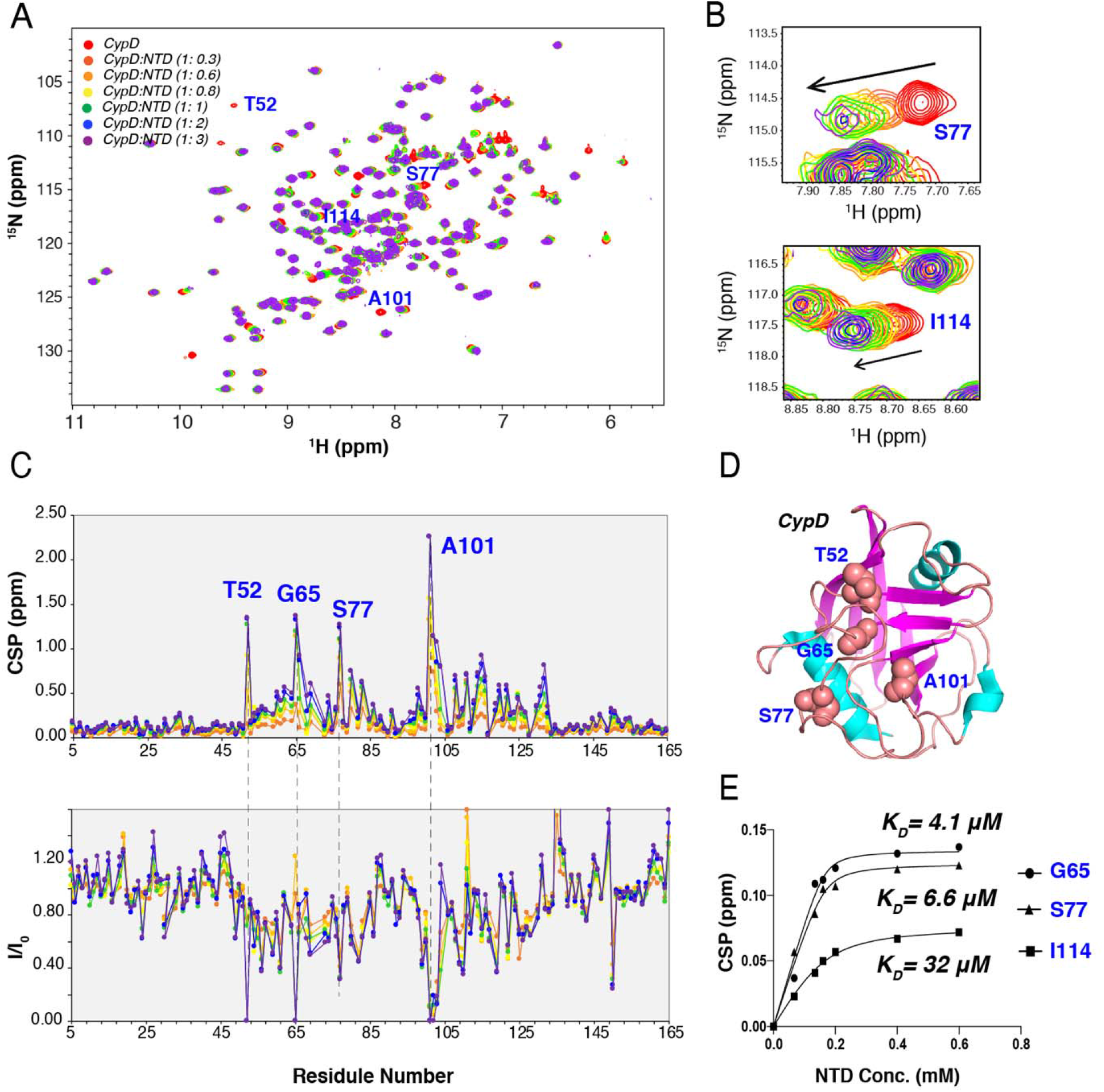
NTD binds CypD with μΜ affinity and an extensive interface. (A) ^1^H-^15^N HSQC spectra of CypD titrated by NTD (ratio 1: 0.3, 1:0.6, 1:0.8, 1:1, 1:2, and 1:3). (B) Peak movement of residue S77 and I114 during titration. (C) CSPs and peak intensity changes (I/I_0_) of CypD titrated by NTD. (D) Residues with largest CSPs were mapped on the crystal structure of CypD (PDB: 3r4g). CypD is shown as a cartoon with α-helix colored in magenta, β-sheet colored in cyan and unstructured region colored in pink. Dominant residues T52, G65, S77 and A101 in the interface are shown as sphere. (E) Residue-based binding affinities (Κ_*D*_) were calculated for residue G65, S77 and I114 by CSP of proton in NMR titration. More residue-based Κ_*D*_ can be found in Fig. S1.

### NTD binds to the CsA-binding face of CypD

As NMR CSPs revealed that NTD can model FLp53 binding to CypD, we carried out a 2D HSQC NMR titration on ^15^N CypD by gradually adding NTD. HSQC spectra of CypD alone and with a CypD: NTD ratio of 1:0.3, 1:0.6, 1:0.8, 1:1, 1:2, and 1:3 were recorded (Fig. 3A) with CSPs and peak intensity changes (I/I_0_) calculated (Fig. 3C). Significant CSPs and intensity decreases were observed for multiple CypD residues with the addition of NTD, with residues A101, S77, G65 and T52 exhibiting the largest CSPs and smallest I/I_0_. After mapping these residues onto the 3D structure of CypD (Fig. 3D), a broad interface was observed, localized to the flexible loop region of CypD. This broad interface is located at the CsA binding face of CypD, also predicted by previous molecular dynamic simulations^25–27^. Adding NTD into the ^15^N-CypD/CsA complex validated this result, which shows no further peak movements, indicating a shared interface by NTD and CsA (Fig S1A). Adding CsA into the ^15^N-NTD/CypD complex showed that CsA reverses the NTD peak movement and disappearance caused by CypD, again supporting a shared interface (Fig S1B). These results agree with previous studies showing that CsA can block p53-CypD interaction^5,18^. Binding affinities for specific residues were calculated by fitting the CSPs in the proton dimension plotted against NTD concentration^28^. Residue G65, S77 and I114 exhibit a binding affinity of 4.1 μΜ, 6.6 μΜ, and 32 μΜ, respectively (Fig. 3E). These residue-based binding affinities are consistent with the NTD-CypD binding affinity measured by SPR (Fig. 1D and 1E).

#### NTD-CypD interaction involves a broad and dynamic interface dominated by electrostatic interactions

To further characterize the molecular basis of p53-NTD-CypD interaction, we performed molecular dynamics (MD) simulations using a Hybrid Resolution (HyRes) protein model designed specifically for modeling intrinsically disordered proteins (see Methods for details). The HyRes model was first verified to provide a realistic description of the conformational properties of unbound p53-NTD compared to previous atomistic simulations and NMR studies (Fig. S4). We then performed 6 groups of independent 500-ns MD simulations with p53-NTD placed 50 Å away from CypD in different directions (Fig. S5). The results showed that p53-NTD binds to the same interface of CypD regardless of the initial placement (e.g., see Fig. S6), giving rise to similar final probabilities of CypD residues involved in contacting p53-NTD (Fig. S7). Analysis of the conformational properties of p53-NTD in the bound state are also consistent from all six groups of independent binding simulations (Fig. S8). Importantly, the simulation correctly predicted that p53-NTD binds to the same CsA binding face of CypD as identified by NMR (highlighted in purple in Fig. S6 and Fig. 4). Together, these results suggest that HyRes is appropriate for characterizing molecular details of the p53-NTD-CypD interactions.

**Fig. 4.**
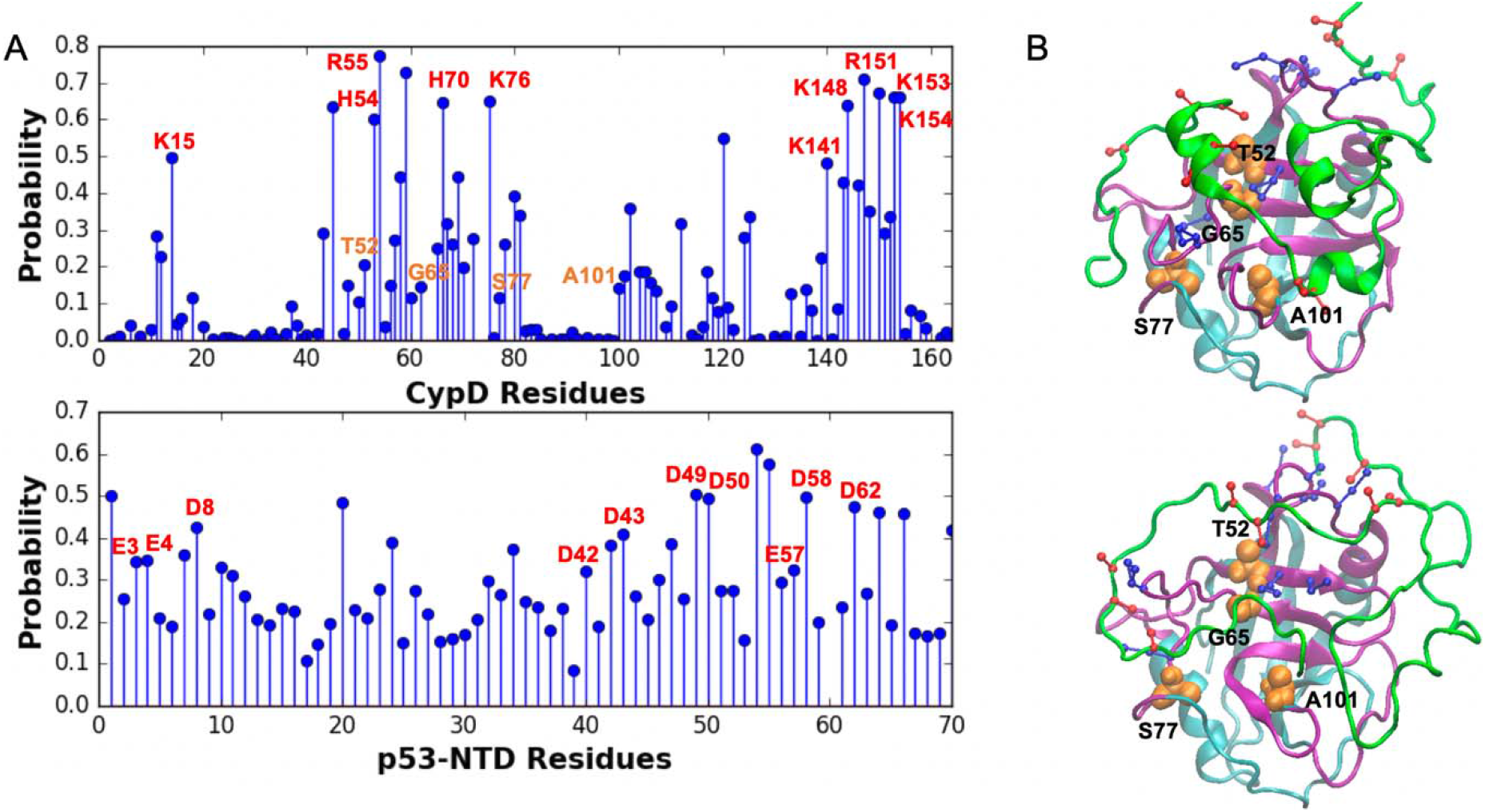
Dynamic interaction between p53-NTD and CypD from molecular modeling. (A) Probability of residues involved in protein-protein interactions. Charged residues are enriched in residues with high probabilities, which predominately involves positively charged residues on CypD and negatively charged residues on p53-NTD. Charged residues with >0.3 probabilities are labelled and shown in sticks in panel B. (B) Two representative snapshots of the bound state. p53-NTD are shown in green cartoon, and CypD in purple (binding interface) and cyan (rest) cartoons. Charged residues labelled in (A) are shown in sticks (red for negative and blue for positive). In addition, four residues (T52, G65, S77 and A101) with the largest NMR CSP values are shown in orange spheres, which locate in the middle of the broad interface. Note that p53-NTD interacts dynamically with CypD, with fluctuating secondary structures and overall conformations.

Further analysis of the simulated conformational ensemble of the complex revealed that the interaction is highly dynamic. The average contact probabilities derived from the last 400 ns of all binding simulation trajectories, shown in Fig. 4A, showed that numerous p53-NTD and CypD residues are involved in transient inter-protein contacts. Furthermore, contacting residues are dominated by charged residues. The binding interface of CypD is greatly enriched with basic residues (highlighted in blue sticks in Fig. 4B), which complement many acidic residues on p53-NTD (highlighted in red sticks in Fig. 4B). This suggests that the p53-NTD and CyD interaction is mainly driven by electrostatic interactions, which also explains the highly dynamic nature. This is not necessarily surprising as NTD has a net charge of −17 and CypD is positively charged with a pK_a_ of 9.2. The dynamic nature of the interaction complicates the interpretation of backbone CSP measurements summarized in Fig. 4, as transient interactions involving charged side chains may or may not significantly perturb the chemical shifts of backbone atoms of the same residue. For example, residue G65 is one of the residues showing the larges CSP values; yet is largely buried and its large CSP does not suggest its direct involvement in inter-protein interactions. Instead, its location near the center of the large binding interface (Fig. 4B) likely allow its backbone chemical shifts to be perturbed by numerous transient interactions nearby. In contrast to a relatively localized binding interface on CypD, the simulation suggests that the whole p53-NTD is engaged in binding, with a slightly elevated probability in the C-terminal D2 region. This is consistent with NMR experiments showing [we should show CSP results at 1:1 or high ratio]. The NTD of p53 remains disordered in the bound state with CypD, as illustrated in Fig. 4B. The average residue helicity of p53-NTD is modestly perturbed in the bound state, and the peptide becomes more extended (Fig. S9).

### p53-CypD interaction is driven by electrostatic interaction

To test the role of electrostatic interaction in p53-CypD interaction revealed in molecular modeling, binding buffers with different NaCl concentrations (150 mM, 250 mM, and 500 mM) were used in SPR experiments for both FLp53 and NTD 1-70. As shown in Fig. 5A, moderate salt concentration (250 mM NaCl) inhibited more than 50% of the binding for both FLp53 and NTD 1-70, compared to control (150 mM NaCl). High salt concentration (500 mM NaCl) almost completely suppressed p53 binding to CypD. SPR measurements of NTD 1-70:CypD interaction showed a much faster dissociation process at 250 mM NaCl, suggesting a less stable complex formed at high salt environment (Fig. 6B and 6C). These results confirmed that the p53-CypD interaction is mainly driven by electrostatic interactions.

**Fig. 5.**
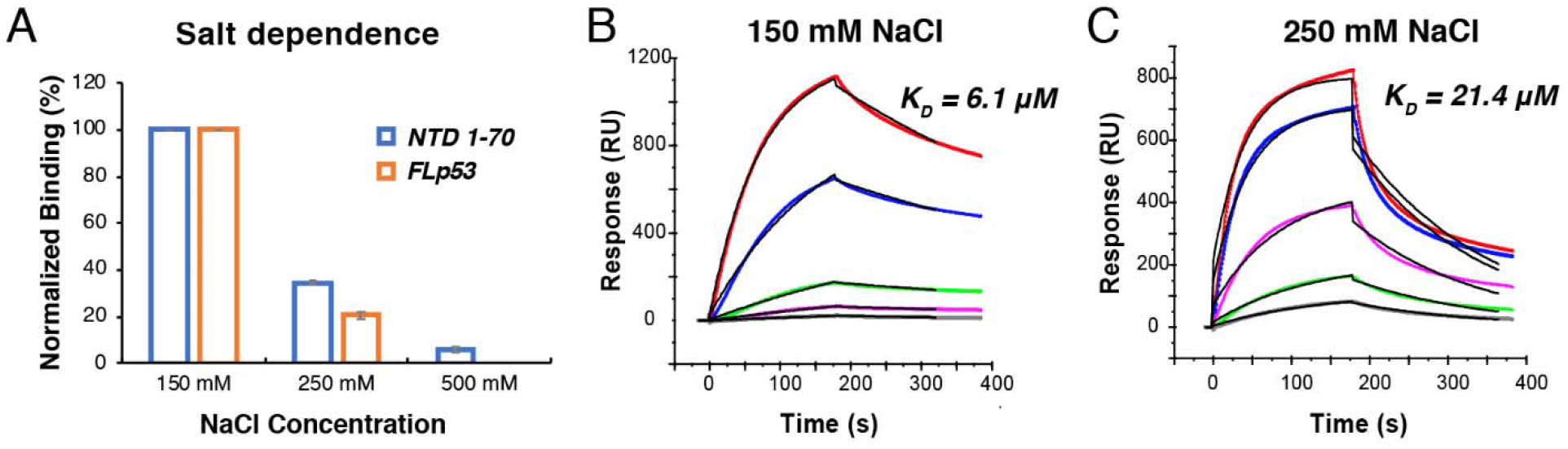
Salt dependence of p53-CypD interaction by SPR. (A) Normalized binding (RU_max_) of FLp53 and NTD1-70 to CypD in SPR. 150 mM NaCl was used as a control. NTD 1-70 was flowed over the chip at 10 μM and FLp53 was at 1μΜ. (B) SPR sensorgrams of the NTD 1-70/CypD interaction at 150 mM NaCl. The concentrations of NTD 1-70 (from top to bottom) were 10, 5, 2.5, 1.25, and 0.63 μM, respectively. (C) SPR sensorgrams of the NTD 1-70/CypD interaction at 250 mM NaCl. The concentrations of NTD 1-70 (from top to bottom) were 30, 20, 10, 5, and 2.5 μM, respectively. All SPR sensorgrams are fitted using 1:1 Langmuir binding model (Black curves) in BIAevaluation software 4.0.1.

**Fig. 6.**
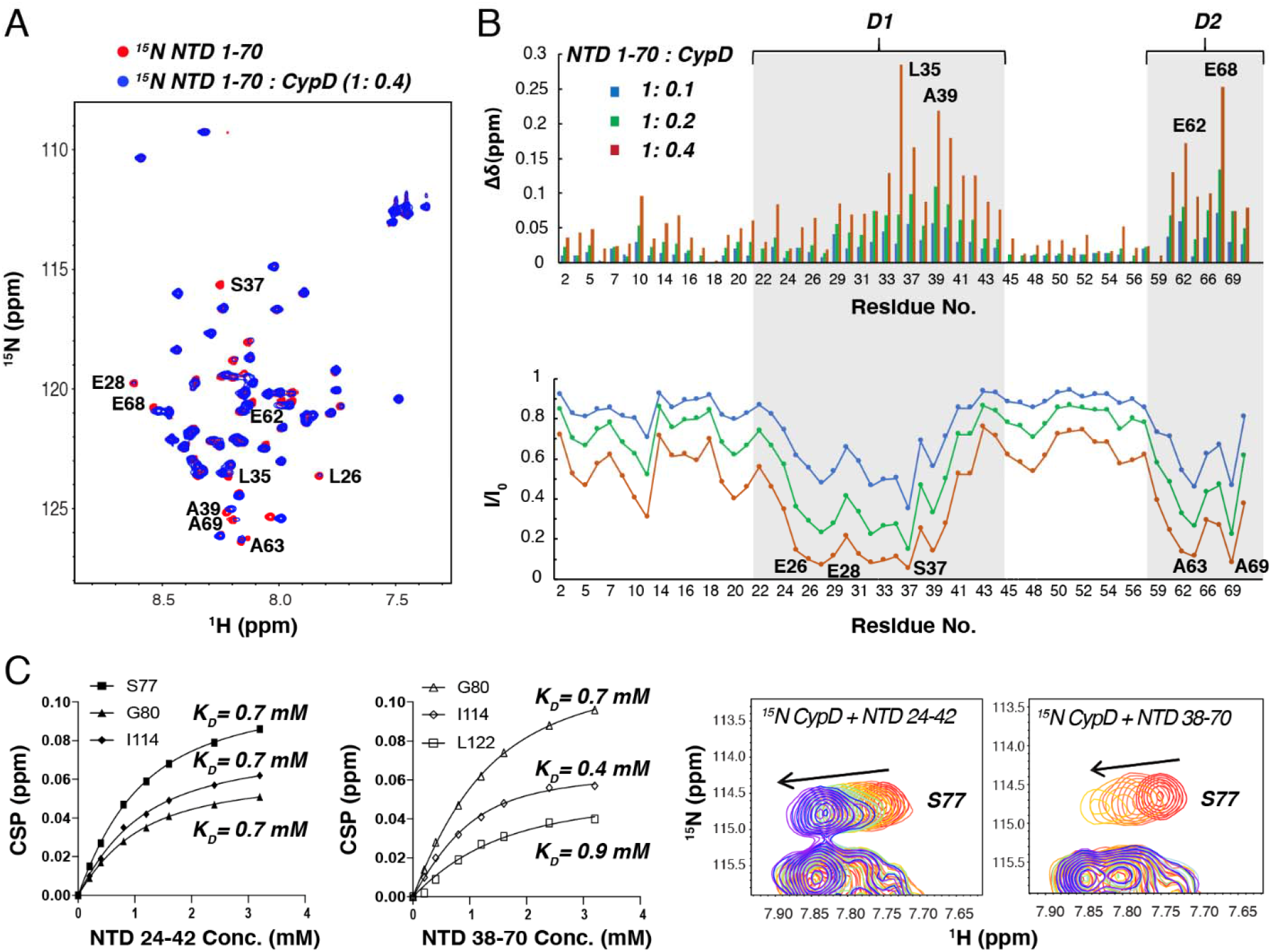
CypD binds NTD at two regions, D1 and D2, both at ~mM affinity. (A) ^1^H-^15^N HSQC spectra of NTD 1-70 upon the addition of CypD (1: 0.4). Peaks with largest CSPs or smallest I/I_0_ were indicated. (B) CSPs and peak intensity changes (I/I_0_) of NTD1-70 titrated by CypD. CypD binding region D1 and D2 were highlighted. (C) NTD24-42 and NTD38-70 bind CypD both at ~mM affinity while different patterns of binding were observed (residue S77 was shown as an example and the full spectra can be referred in Fig. S3).

### Two NTD fragments bind CypD with mM affinity

To thoroughly understand the binding interface, NMR experiments were performed on isotopically labeled NTD with the addition of CypD. Since NTD 1-70 showed comparable binding affinity to CypD with full-length NTD (Fig.1D and 1E), an NMR titration experiment was carried out at low NTD 1-70: CypD ratios (NTD: CypD ratio of 1: 0.1, 1: 0.2 and 1: 0.4). As shown in Fig. 6A, NTD 1-70 retains the intrinsically disordered structure of full length NTD indicated by the pattern of HSQC spectrum. Upon the addition of CypD (1: 0.4), HSQC spectrum of NTD1-70 experienced significant peak movement and/or intensity changes. Residues with largest CSPs include L35, A39, E68, and E62, which were located in two different regions on NTD 1-70, which we named as D1 (residues 22-44) and D2 (residues 59-70) (Fig. 6B). Residues with the largest intensity changes were also located in D1 and D2, corresponding to S37, E28, E26 in D1 and A63, A69 in D2. Interestingly, residues like E26 and E28, which show minimal CSPs, exhibit dramatical peak intensity decrease, indicating complex binding dynamics in D1. To test if both D1 or D2 alone can bind CypD, two peptides, NTD 24-42 and NTD 38-70 were synthesized and their binding to CypD were evaluated by SPR and NMR titration. In contrast to NTD 1-70, 10 μΜ of NTD 24-42 and NTD 38-70 showed no sign of binding associated RU increase in SPR, indicating that NTD 24-42 and NTD 38-70 bind CypD at much lower affinity (Fig. S2).

Their binding affinities were measured to be in the mM range by NMR titration (Fig. 6C), consistent with minimal binding in SPR. These data suggest that D1 or D2 alone is not sufficient for μM affinity CypD binding. Although NTD 24-42 and NTD 38-70 have similar binding affinity to CypD, different binding behaviors were observed for two peptides at certain regions. For example, only CSP was observed for residue S77 upon titration of NTD 24-42; while its peak intensity decreased greatly when titrated by NTD 38-70 (Fig. 6C). In addition, upon titration of NTD 24-42, CypD residues around R55 show more significant CSPs (Fig. S3A and S3C); while more peak intensity decreases were observed upon titration of NTD38-70 (Fig. S3B and S3D), again implying that NTD domain D1 and D2 may play different roles in the binding of CypD.

### NTD 20-70 mimics NTD binding to CypD in both SPR and NMR

As both NTD region D1 and D2 are required for high affinity CypD interaction, a peptide NTD 20-70 was overexpressed and purified to characterize its binding to CypD. CSPs generated by NTD 20-70 in CypD resemble those caused by full-length NTD, indicating NTD 20-70 binds CypD in the same manner with full-length NTD (Fig. 7A). Consistently, a binding affinity *K_D_* of 9.4 μM was obtained for CypD/NTD 20-70 interaction from SPR (Fig. 7B), close to that of 8.4 μM for CypD/NTD interaction (Fig. 1D).

**Fig. 7.**
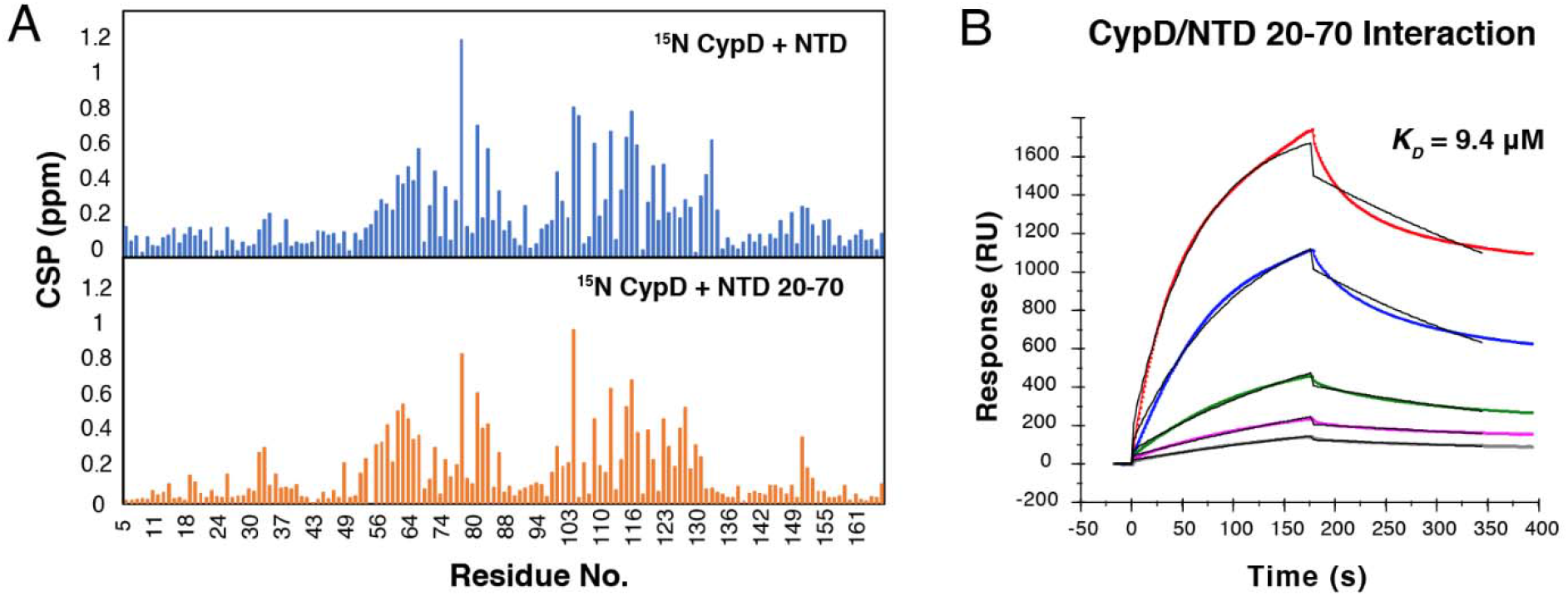
NTD 20-70 mimics full-length NTD binding to CypD with μΜ affinity. (A) Addition of NTD 20-70 into ^15^N CypD resembles the CSPs of CypD caused by full-length NTD. Molar ratio for CypD and NTD/NTD20-70 was 1:2. (B) SPR shows an affinity *K_D_* of 9.4 μM for CypD/NTD 20-70 interaction. The concentrations of NTD20-70 (from top to bottom) were 40, 20, 10, 5, and 2.5 μM, respectively. SPR sensorgrams are fitted using 1:1 Langmuir binding model (Black curves) in BIAevaluation software 4.0.1.

## Discussion

While mitochondrial translocation of p53 and its phenotype are established to modulate mitochondrial permeability through interaction with CypD, the molecular details of p53/CypD interaction are not well understood. In this study, we identify the intrinsically disordered N-terminal domain (NTD) of p53 as the major contributor to the p53-CypD interaction using SPR, NMR and MD. A ~μΜ binding affinity was obtained for NTD-CypD interaction, which is similar to that of FLp53 and CypD. NTD binds CypD at a broad surface on CypD, overlapping with the CsA binding interface. The interaction is highly dynamic and dominated by electrostatic interactions. Two CypD-binding sub-domains D1 and D2 with mM affinity were mapped on NTD. NTD 20-70 containing both D1 and D2 was identified as the minimal binding region to maintain μΜ binding with CypD.

Notably, discrepancies were found between our results and previous studies. Based on immunoprecipitation assays, Moll et al. concluded that p53 residues 80–220, corresponding to the N-terminal portion of the DBD, is required for CypD binding. In a followed-up study, Seeliger and Moll et al. mapped the binding of DBD on CypD by NMR titration ([CypD]: [DBD] = 1:2). However, overall, very small perturbations were observed on CypD upon DBD binding, indicating mM or lower affinity. In contrast, much larger CSPs were observed on CypD upon titration of NTD in this study (Fig. 3), suggesting a much higher affinity, which was confirmed by both SPR kinetic measurements (Fig. 1) and NMR-based affinities (Fig. 3E). Our data clearly demonstrated that NTD plays a critical role for CypD binding, rather than DBD. C-terminal truncated isoforms of p53, including naturally occurring p53Ψ (lacking the major portions of DBD) and exon-6 truncating mutations (R196* and R213*; lacking NLS) have been found to translocate into mitochondrial. These p53 variants increase mPTP opening through CypD binding, supporting a role of p53 N-terminus in the CypD interaction and mitochondrial permeability modulation.

In summary, we report for the first time that CypD interacts with the NTD of p53, with extensive characterization of the molecular details in NTD-CypD interaction and validated the idea that FLp53 interacts with CypD mainly through the N-terminal domain. Our work provides novel molecular insights for the p53-CypD interaction and has important implications on the p53-dependent mPTP opening and apoptosis, and drug discovery efforts targeting p53-CypD interaction.

## Method

### Materials

Full-length p53, NTD, NTD-DBD, NTD1-70 and NTD20-70 were overexpressed and purified as previously described^29^. Briefly, p53 coding sequences were inserted downstream of the *Thermoanaerobactor tencongenesis* ribose binding protein (RBP) gene, fused with a nucleotide sequence that harbors an HRV3C protease cleavage site. A His-tag sequence was placed at the 3’ end of the NTD gene to facilitate purification of the full-length fusion protein. The fusion gene was placed in the pET41 plasmid (Novagen) and transformed into *E. coli* BL21(DE3). Proteins were expressed at 18°C with induction by 20 mg/L of isopropyl β-D-1-thiogalactopyranoside (IPTG) and purified on a Ni^2+^-NTA column. Following overnight cleavage by HRV3C protease (4 °C), the NTD proteins were recovered by Ni^2+^-NTA and S75 size exclusion chromatography to a final purity of >95% as judged by SDS-PAGE. The NTD constructs contain GPG and LE(H)_8_ sequences at the N- and C-termini, respectively. Peptides NTD 24-42 and NTD 38-70 were synthesized by Genscript (NJ, USA). CypD was overexpressed and purified following protocols from Schlatter et al.^30^. Briefly, the overexpression plasmid was transformed into *E. coli* BL21(DE3). Protein was expressed at 30°C and induced with 0.1 mM IPTG. Cells were collected and the protein was purified by applied to a HI-PREP SP 16/10 cation exchange column and a HI-PREP Q 16/10 anion exchange column from GE Healthcare Biosciences (Pittsburgh, PA, USA) on a FPLC from GE Healthcare Biosciences (Pittsburgh, PA, USA). ^15^N labeled samples were achieved by growing cells in 1 g/L of ^15^NH_4_Cl and 4 g/L of glucose. ^15^N and ^13^C doubly labeled samples were obtained by growing cells in 1 g/L of ^15^NH_4_Cl and 4 g/L of ^13^C_6_-D-glucose (Cambridge Isotope Laboratories, MA, USA). SPR CM5 sensor chip and amine coupling kits were obtained from GE Healthcare Bio-Sciences AB (Uppsala, Sweden).

### p53-CypD interaction by SPR

#### Binding kinetics of CypD and different p53 constructs

The binding behavior of different p53 constructs with CypD was measured using surface plasmon resonance (SPR) on a BIAcore 3000 system. First, CypD was immobilized on a CM5 chip through its amine groups using EDC/NHS according to the standard amine coupling protocol. A 10 μL solution of 0.1 mg/mL CypD was injected over the flow cell at 5 μL /min. Successful immobilization of CypD was confirmed by an ~3000 resonance unit (RU) increase in the sensor chip. The first flow cell (control) was prepared without injection of CypD. After immobilization, different p53 samples were diluted in running buffer (20 mM Tris-HCl, 150 mM NaCl, 1 mM TCEP, pH7.2), separately. Different dilutions of p53 were injected at a flow rate of 30 μL/min for 3 minutes. Following sample injection, running buffer was passed over the sensor surface for a 3-minute period for dissociation. The sensor surface was regenerated by a 30 μL injection of 2M NaCl. The response was determined as a function of time (sensorgram) at 25°C. Binding kinetics were recorded and the binding affinity (*K_D_*) was calculated by fitting to a 1:1 Langmuir binding model in BIAevaluation software 4.0.1 (GE Healthcare).

#### Salt dependence

FLp53 and NTD 1-70 were diluted in HBS-P buffer (0.01 M HEPES, 0.15 M NaCl, and 0.005% surfactant P20, pH 7.4) with different NaCl concentrations (150 mM, 250 mM, and 500 mM) and injected at a flow rate of 30 μL/min. At the end of the sample injection, the same buffer was flowed over the sensor surface to facilitate dissociation. After a 3 min dissociation time, the sensor surface was regenerated by injecting 30 μL of 2 M NaCl. The response was monitored as a function of time (sensorgram) at 25°C. Binding kinetics of CypD and NTD 1-70 in the presence of 150 mM NaCl and 250 mM NaCl were recorded by injecting a series of NTD 1-70 dilutions and the binding affinity (*K_D_*) was calculated by fitting to a 1:1 Langmuir binding model in BIAevaluation software 4.0.1 (GE Healthcare).

### 2D NMR on ^15^N CypD

2D ^1^H-^15^N heteronuclear single quantum coherence (HSQC) NMR spectra of CypD were acquired at 25°C on a Bruker 800 MHz NMR spectrometer equipped with a cryogenic probe. NMR data were processed and analyzed using Topspin 3.2pl7 and Sparky 3.115. ^15^N-labled CypD was dissolved in 100 mM NaCl, 50 mM Na_2_HPO_4_, 1 mM EDTA and 2 mM DTT at pH 7.2 in 90/10% H_2_O/D_2_O. Chemical shift perturbation (CSP) in the amide nitrogen (N) and amide proton (H) dimension upon FLp53/ NTD/ NTD-DBD/ NTD 1-70/ NTD 20-70/ NTD 24-42/ NTD 38-70 binding were calculated by the difference between chemical shifts of the free and bound form of CypD, represented by Δ*N* and Δ*H*, respectively. The normalized, weighted average CSP for amide ^1^H and ^15^N chemical shifts upon p53 addition were calculated using the equation 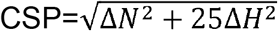. For NTD, a series of HSQC experiments were performed on a 0.1 mM ^15^N CypD sample by adding increasing amounts of p53 aliquots. Residue-based binding affinity was calculated using Δ*H* by equation ^28^ Δ*H*_obs_ = Δ*H*_max_ {([P]_t_ + [L]_t_ + K_D_-([P]_t_ + [L]_t_ + K)^2^ - 4[P]_t_ [L]_t_ ^1/2^}/2[P]_t_, where Δ*H*_max_ is the maximum shift change on saturation, [P]_t_ is the total CypD concentration, [L]_t_ is the total NTD concentration, and K_D_ is binding affinity.

### 2D NMR on ^15^N NTD

2D ^1^H-^15^N HSQC NMR spectra of NTD 1-70 were acquired at 25°C on a Bruker 800 MHz NMR spectrometer equipped with a cryogenic probe. NMR data were processed and analyzed using Topspin 3.2pl7 and Sparky 3.115. ^15^N-labled NTD 1-70 was dissolved in 50 mM NaCl, 25 mM Na_2_HPO_4_, 1 mM EDTA and 1 mM DTT at pH 6.9 in 90/10% H_2_O/D_2_O. Chemical shift perturbation (CSP) in the amide nitrogen (N) and amide proton (H) dimension upon CypD binding were calculated by the difference between chemical shifts of the free and bound form of CypD, represented by Δ*N* and Δ*H*, respectively. The normalized, weighted average CSP for amide ^1^H and ^15^N chemical shifts upon p53 addition were calculated using the equation 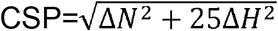.

### Molecular modeling of NTD and its interaction with CypD

#### Simulation protocol and benchmarks

A multiscale hybrid resolution (HyRes) protein model^31^ was used to model the dynamic interaction between p53-NTD and CypD. The HyRes model contains atomistic representation of the protein backbone, to provide semi-quantitative secondary structure propensities, while the sidechains are described in an intermediate resolution, to provide a qualitative description of long-range nonspecific interactions. The appropriateness of using the HyRes model was first evaluated by simulation of the unbound p53-NTD. For this, the N- and C- termini of the p53-NTD peptide (residues 1-70) were capped with an CH_3_CO- group and CH_3_NH-, respectively. A fully extended conformation was then generated to initiate two 1000 ns simulations at 300 K using CHARMM^32,33^. Default HyRes options were used for non-bonded interactions^31^. Langevin dynamics was always used with a friction coefficient of 0.2 ps^−1^ and a time step of 2 fs. The resulting ensembles are well-converged and successfully recapitulate the average helicity derived from previous atomistic simulations and NMR^34^ (Fig. S4).

For the simulation of the p53-NTD-CypD interaction, the atomistic structure of CypD^3^ (PDB: 2z6w) was first mapped on to the HyRes model (Fig. S5A) and then minimized with Cα atoms harmonically restrained. One pair of p53-NTD and CypD molecules was placed in the center of the simulation box, with p53-NTD initially placed around 50 Å away in 6 directions (Fig. S5B). The initial structure of p53-NTD was extracted from the equilibrium simulation of p53-NTD after its convergence (see Fig. S4). The complex was simulated at 300 K and the backbone atoms of CypD was restrained by harmonic potentials with a force constant of 1.0 kcal/mol to prevent structural deformation during simulation. For each of the 6 initial placements, four parallel simulations were performed with the initial p53-NTD conformation randomly rotated. Periodic boundary conditions were imposed with a cubic box size 120 Å. The total simulation time for each initial condition (6×4) was 500 ns, which was found to be adequate for sampling the dynamic interactions between two proteins.

#### Analysis of simulation trajectories

All structural analyses were performed using CHARMM and in-house scripts unless otherwise specified. Contacts between two residues are considered formed if the minimum distance is no greater than 5 Å. The first 100 ns of all trajectories was considered equilibration and not included in calculation of probability distributions. Convergence of the simulations was evaluated by comparing various distributions derived from independent simulations, s1-2 for unbound (Fig. S4) and s1-6 for bound states (Fig. S6-9). The results for each set of the complex simulation were derived from averaging over four parallel simulations with different initial random rotation of p53-NTD (Fig S5). All molecular illustration was prepared using VMD^35^.

## Acknowledgements

Please include the following: The authors thank Dr. Xiaorong Liu for help with the HyRes model. Grant support: NIH R01-GM114300 to J. C

## Notes

### Competing Interest Statement

The authors have declared no competing interest.

### Summary of Updates

Author order revised.

