## Supplemental Figures for "Intrinsically disordered N-terminal domain (NTD) of p53 interacts with mitochondrial PTP regulator Cyclophilin D" (from another version of this preprint: version 1, 2021-07-23)

**Supplementary Information**

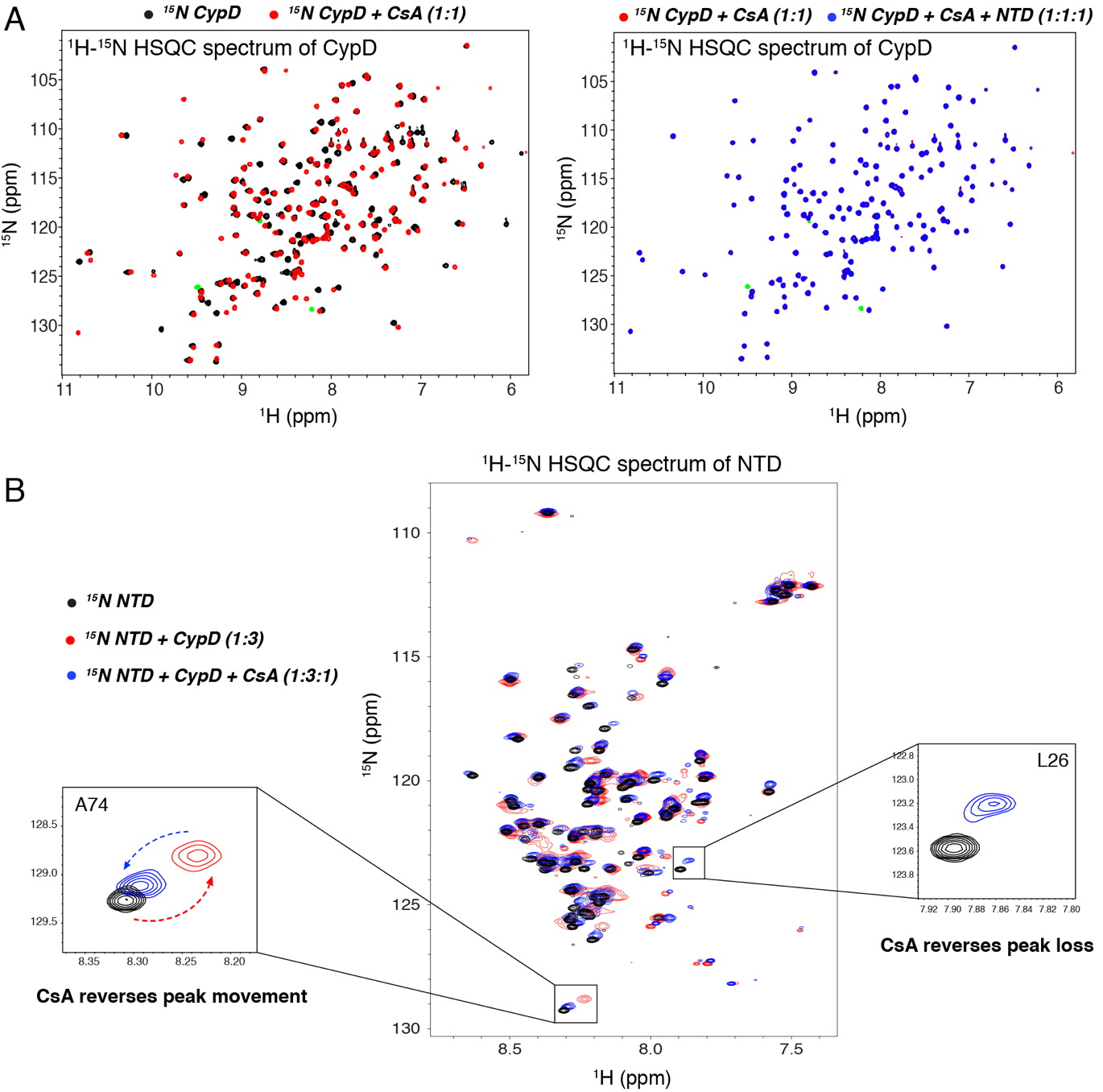

**Fig. S1.** Competitive NMR validates NTD binds to the CsA binding face of CypD. (A) Adding NTD into the ^15^N-CypD/CsA complex (1:1) shows no further peak movements. The concentration of ^15^N-CypD was 100 μΜ. (B) Adding CsA into the ^15^N-NTD/CypD complex (1:3) showed that CsA reverses the NTD peak movement and disappearance caused by CypD. The concentration of ^15^N-NTD was 300 μΜ.

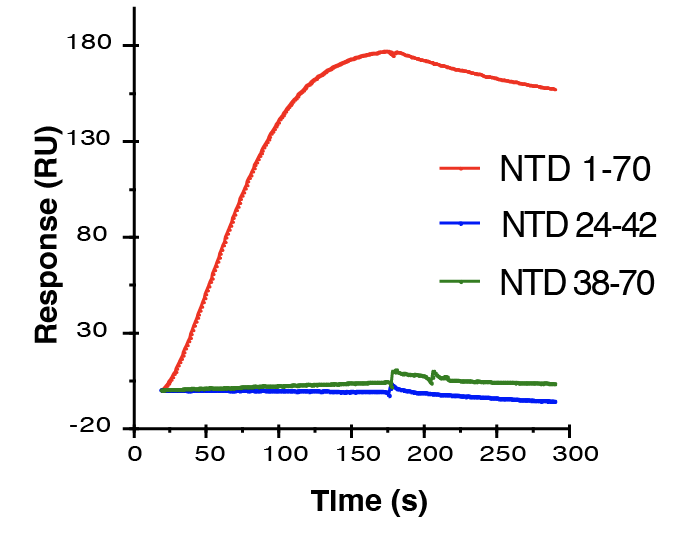

**Fig. S2.** Minimal binding of CypD with two peptides NTD24-42 and NTD38-70 in SPR. The concentration of NTD1-70/NTD24-42/NTD38-70 was 10 μM.

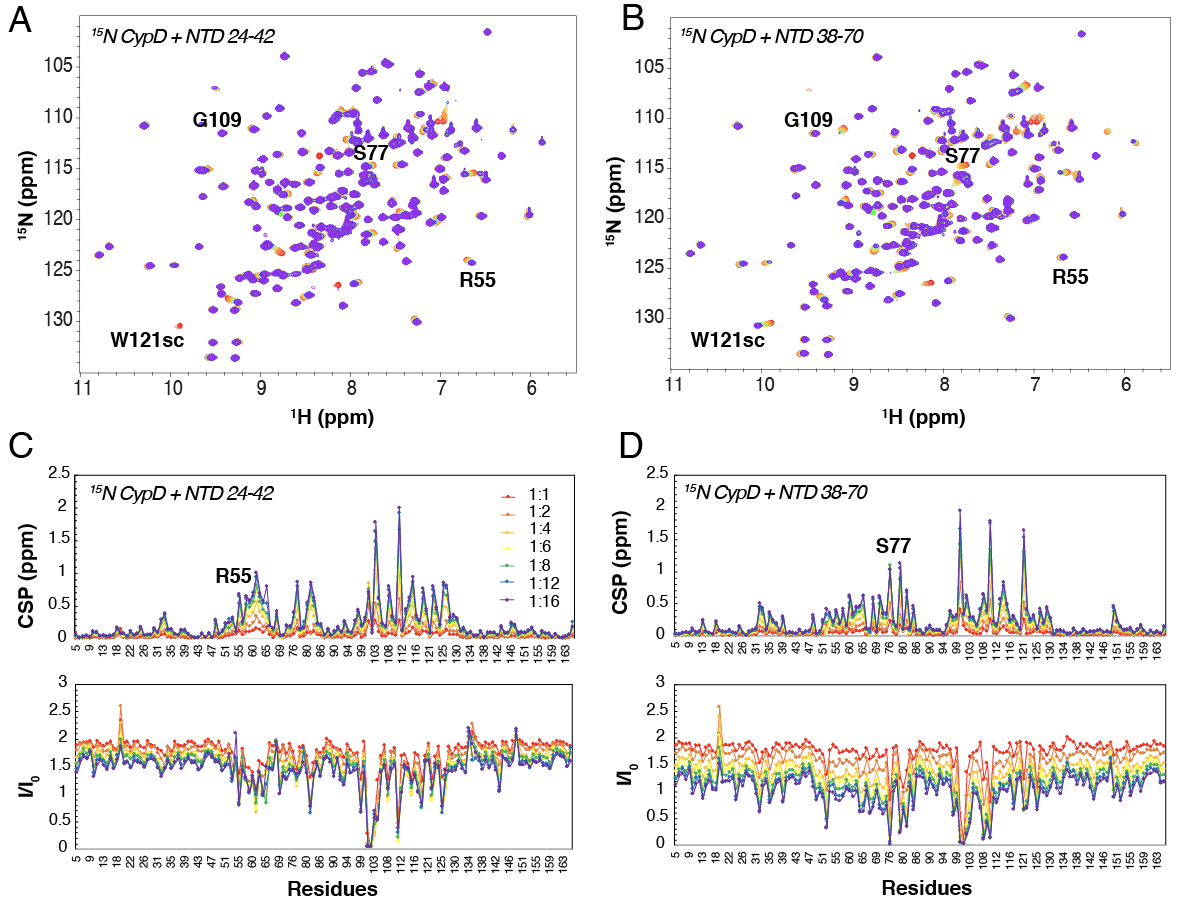
**Fig. S3.** NMR titration of NTD 24-42 and NTD 38-70 into ^15^N CypD. (A) ^1^H-^15^N HSQC spectra of CypD titrated by NTD 24-42 (ratio 1:1, 1:2, 1:4, 1:6, 1:8, 1:12 and 1:16). (B) ^1^H-^15^N HSQC spectra of CypD titrated by NTD 38-70 (ratio 1:1, 1:2, 1:4, 1:6, 1:8, 1:12 and 1:16). (C) CSPs and peak intensity changes (I/I_0_) of CypD titrated by NTD 24-42. (D) CSPs and peak intensity changes (I/I_0_) of CypD titrated by NTD 38-70.

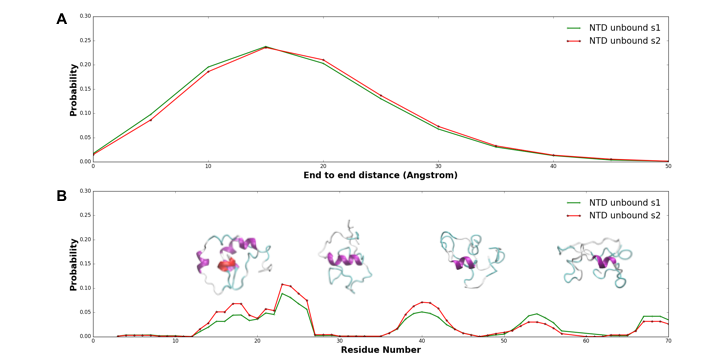

**Fig. S4.** Convergence and conformational properties of p53-NTD in the HyRes model. (A) End-to-end distance and (B) Average residue helicity profile. The results were calculated from two independent 1-μ simulations, s1 (green) and s2 (red). Four representative snapshots are also shown, with the helical segments colored in purple.

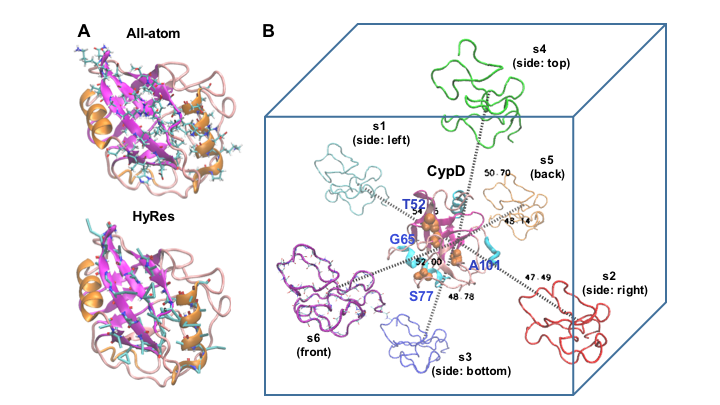

**Fig. S5.** A) The structure of CypD in atomistic and HyRes representations. Note that HyRes contains atomistic backbone and intermediate side chains. B) Initial configurations of binding simulations, with CypD placed at the central of the periodic box and p53-NTD 50 Å away in six different orientations. The orientation of p53-NTD was randomly rotated prior to each independent simulation.

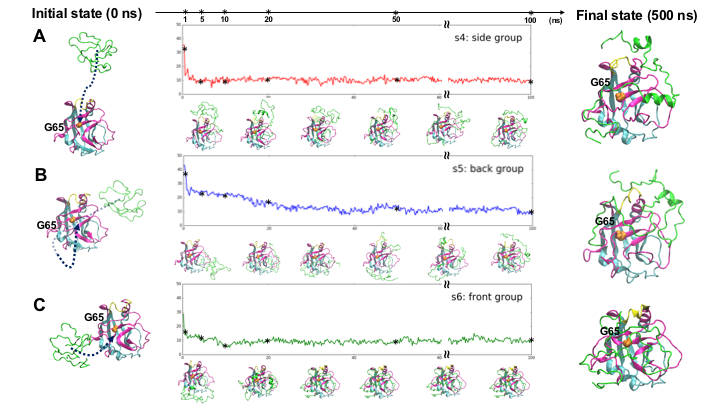

**Fig. S6.** Representative trajectories of p53-NTD binding to CypD with p53-NTD initial placed at either the side (A), back (B) and front (C) of the final binding interface. The traces show the evolution of minimum distance from p53-NTD to CypD G65, which locates around the center of the binding interface. Snapshots are shown at initial, 1 ns, 5 ns, 10 ns, 20 ns, 50 ns, 100 ns and finial (500 ns) timepoints for each trajectory to illustrate the evolution of the interaction interface. The approximate paths of p53-NTD binding to CypD are shown as blue dashed lines in the initial state snapshots. In all snapshots, p53-NTD is shown as green cartoon and CypD in purple/yellow (binding interface) and cyan (rest of the protein) cartoons.

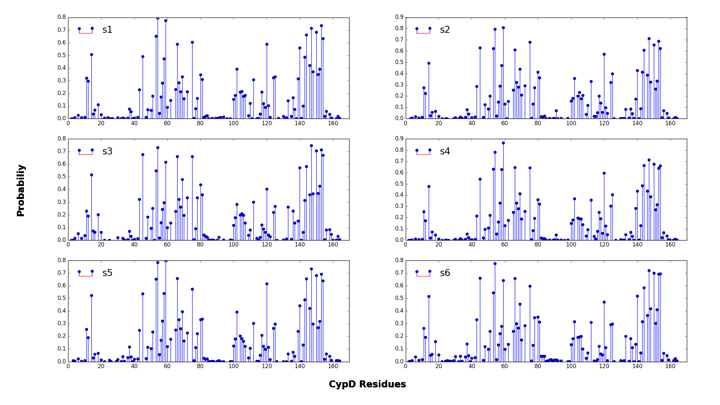

**Fig. S7.** Probabilities of CypD residues involved in contacting p53-NTD derived from six set of 500-ns MD simulations (s1-6) with p53-NTD placed in different initial directions (Fig. S5). Each set’s result was averaged from the bound conformations sampled during the last 500 ns of four independent simulations.

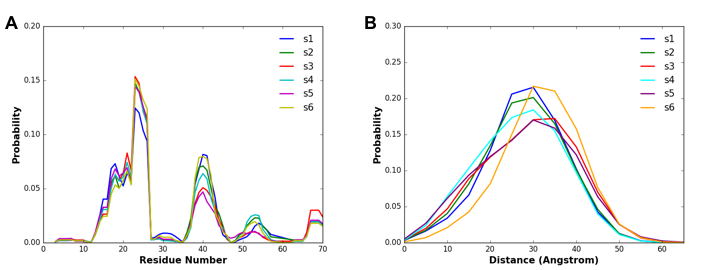

**Fig. S8.** Conformational properties of p53-NTD in bound state derived from 6 sets of independent simulations (see Fig. S5). (A) Average residue helicity profiles and (B) Distributions of the end-to-end distance. The result of each set was averaged from the bound conformations sampled during the last 500 ns of four independent simulations.

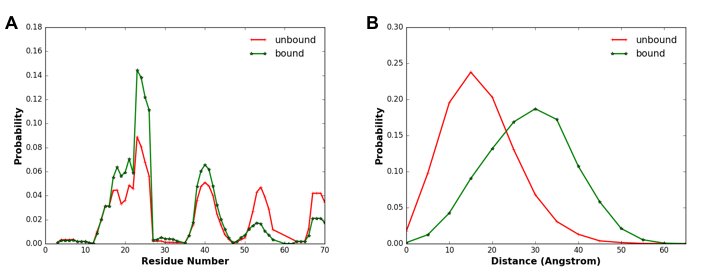

**Fig. S9.** Conformational properties of p53-NTD in unbound (red) and bound (green) states. (A) Average residue helicity profiles and (B) Distributions of the end-to-end distance. The results were calculated from all simulation trajectories of the unbound p53-NTD and p53-NTD/CypD complex.
